# Lifestyle-intervention-induced reduction of abdominal fat is reflected by a decreased circulating glycerol level and an increased HDL diameter

**DOI:** 10.1101/718023

**Authors:** Marian Beekman, Bianca A.M. Schutte, Erik B. van den Akker, Raymond Noordam, Petra Dibbets-Schneider, Lioe-Fee de Geus-Oei, Joris Deelen, Ondine van de Rest, Diana van Heemst, Edith J.M. Feskens, P. Eline Slagboom

## Abstract

Abdominal obesity is one of the main modifiable risk factors of age-related cardiometabolic disease. Cardiometabolic disease risk and its associated high abdominal fat mass, high cholesterol and glucose concentrations can be reduced by a healthier lifestyle. Hence, our aim is to understand the relation between lifestyle-induced changes in body composition, and specifically abdominal fat, and accompanying changes in circulating metabolic biomarkers.

**Methods and results:** We used the data from the Growing Old Together (GOTO) study, in which 164 older adults (mean age 63 years, BMI 23-35 kg/m^2^) changed their lifestyle during 13 weeks by 12.5% caloric restriction plus 12.5% increase in energy expenditure. We show that levels of circulating metabolic biomarkers, even after adjustment for body mass index, specifically associate with abdominal fat mass. Next, we show that the applied lifestyle intervention mainly reduces abdominal fat mass (−2.6%, SD=3.0) and that this reduction, when adjusted for general weight loss, is highly associated with decreased circulating glycerol concentrations and increased HDL diameter.

**Conclusions:** The lifestyle-induced reduction of abdominal fat mass is particularly associated, independent of body mass index or general weight loss, with associated with decreased circulating glycerol concentrations and increased HDL diameter.

## 1. Introduction

Abdominal obesity plays an important role in the development of cardiometabolic disease risk^[1,2]^. People with relatively high amounts of abdominal fat are characterized by increased insulin resistance^[3]^ and a detrimental circulating metabolic biomarker profile encompassing high levels of glucose, cholesterol and triglycerides^[4]^, all of which are known to be associated with type 2 diabetes and cardiovascular disease^[5–12]^. Reduction of cardiovascular risk can be achieved by lifestyle interventions aimed at increasing physical activity and/or reducing caloric intake^[13,14]^. Because abdominal fat is intimately linked to disease risk, it is imperative to understand the relation between the lifestyle-induced changes in body composition, specifically abdominal fat, and the accompanying changes in metabolic biomarkers.

To gain more insight than what would be achieved by only measuring the standard metabolic clinical chemistry parameters, such as cholesterol and glucose levels, state-of the art ^1^H-NMR metabolomics platforms have been used to investigate the relationship between metabolism and body composition. In young people (age between 25 and 30 years), a larger amount of abdominal fat has been associated with an unfavourable lipoprotein profile (i.e. high VLDL, IDL, LDL and small HDL particle concentrations, high IDL- and LDL-cholesterol, triglycerides, ApoB and ApoB to ApoA1 ratio and low large HDL particle concentration, HDL-cholesterol and small HDL diameter)^[5]^. In general, an unhealthy metabolic profile, as measured by the ^1^H-NMR platform, can be improved by a lifestyle change^[13–16]^ that is known to particularly reduce the amount of abdominal fat mass^[17–19]^. However, it remains unclear how and to what extent lifestyle-induced changes in body composition, specifically the reduction in abdominal fat, are reflected by circulating metabolic biomarkers.

We investigated the relation between lifestyle-induced changes in body composition and the altering blood metabolome, by exploring the data collected in the Growing Old Together study (GOTO); a 13-week-lifestyle intervention study in which older participants (N=164, Age_mea_n=63 years old (age range 49-75 years), BMI_mean_=27 (BMI range 23-35 kg/m^2^) at the moment of inclusion) increased physical activity by 12.5% and decreased energy intake by 12.5%^[13]^. Body composition parameters were measured with anthropometrics and a DXA scan, while metabolic biomarkers were measured in serum using ^1^H-NMR metabolomics, both before and after the intervention. First, at baseline we cross-sectionally correlated body composition measures with circulating metabolic biomarkers levels. Second, we determined the effect of the GOTO lifestyle intervention on ^1^H-NMR metabolomic biomarkers. Third, we determined how body composition measures were affected by the lifestyle intervention. Finally, we investigated the associations between the change in multiple measures of body composition and the changes in metabolic biomarkers to determine which of these biomarkers reflected the alterations in body composition by a lifestyle change.

## 2. Experimental section

### 2.1 Study design

The GOTO study has previously been described by van de Rest *et al.*^[13]^. The Medical Ethical Committee of the Leiden University Medical Center approved the study and all participants signed a written informed consent. All experiments were performed in accordance with relevant and approved guidelines and regulations. This trial was registered at the Dutch Trial Register (http://www.trialregister.nl) as NTR3499.

In short, the lifestyle intervention comprised 13 weeks of 25% lowered energy balance by 12.5% reduction in energy intake and 12.5% increase in physical activity under supervision of a dietician and a physiotherapist. Participants were recruited within the Leiden Longevity Study^[20]^, consisting of a member of a long-lived family and its partner. Participants (N=164) were between 46 and 75 years (mean age 63 years), had a BMI between 23 and 35 kg/m^2^ (mean BMI = 27 kg/m^2^), no diabetes (fasting glucose <7.0 mmol/L) or any disease or condition that seriously affects body weight and/or body composition including active types of cancer (Table S1).

The participants provided a report of their pharmacist about their current medication use, from which the use of lipid lowering medication (fibrates, niacin, bile acid sequestrants, 3-hydroxy-3-methylglutaryl-coenzyme A reductase inhibitors) and hypertension medication (diuretics, beta-blockers, calcium channel blockers, agents acting on the renin-angiotensin system) was deduced.

In the present paper the analyses have been performed on the subgroup of 132 participants for whom we had data on anthropometrics, DXA measures and NMR metabolomics (Nightingale Health) were available at baseline as well as at the endpoint of the study (Table 1).

**Table 1.** Baseline characteristics of the GOTO study population*

|  | **N** | **Mean** | **SD** |
| --- | --- | --- | --- |
| Age (years) | 132 | 62.8 | 6.0 |
| % Female | 65 | 49.2 |  |
| % Lipid lowering medication | 23 | 17.4 |  |
| % Antihypertensive medication | 41 | 31.1 |  |
| **Anthropometrics** |  |  |  |
| Height (m) | 132 | 1.71 | 0.09 |
| Weight (kg) | 132 | 79.5 | 10.0 |
| Body mass index (kg/m<sup>2</sup>) | 132 | 27.0 | 2.5 |
| Waist circumference (cm) | 132 | 96.1 | 8.1 |
| Hip circumference (cm) | 132 | 104.2 | 5.2 |
| Waist/hip ratio (cm) | 132 | 0.9 | 0.1 |
| Waist/height ratio | 132 | 56.1 | 4.7 |
| **DXA measures<sup>#</sup>** |  |  |  |
| Whole body lean mass (kg) | 132 | 54.3 | 9.8 |
| Whole body fat (kg) | 132 | 25.6 | 6.2 |
| Whole body fat (%) | 132 | 32.3 | 7.4 |
| Trunk fat (kg) | 132 | 13.1 | 3.6 |
| Trunk fat (%) | 132 | 32.6 | 7.2 |
| Android fat (kg) | 132 | 2.2 | 0.7 |
| Android fat (%) | 132 | 35.0 | 7.3 |
| Gynoid fat (kg) | 132 | 4.1 | 1.1 |
| Gynoid fat (%) | 132 | 32.8 | 8.3 |
| Leg fat (kg) | 132 | 8.3 | 2.7 |
| Leg fat (%) | 132 | 32.2 | 9.5 |
| Trunk fat /whole body fat ratio | 132 | 0.51 | 0.06 |
| Android fat /whole body fat ratio | 132 | 0.09 | 0.02 |
| Gynoid fat /whole body fat ratio | 132 | 0.16 | 0.02 |
| Leg fat /whole body fat ratio | 132 | 0.32 | 0.06 |
| Android fat/gynoid fat ratio | 132 | 1.10 | 0.20 |
| Whole body fat/Whole body lean mass ratio | 132 | 0.49 | 0.17 |

\*The subgroup of 132 participants (out of the 164) having data on anthropometrics, DXA measures and <sup>1</sup>H- NMR metabolomics
<sup>#</sup>See Supporting Information for recognition of body regions in DXA images (Figure S1)
SD= standard deviation

### 2.2 Body composition measurements

We had data available for 7 anthropometric measures based on weight, height, waist circumference and hip circumference. Weight was measured to the nearest 0.1 kg using a digital personal scale (Seca Clara 803 scale, Seca Deutschland, Hamburg, Germany) with the person dressed in light clothing and without shoes. Height, waist circumference (midpoint between the lowest rib and the top of the iliac crest) and hip circumference (largest circumference of buttocks) were measured to the nearest 0.1 cm with a non-elastic tape in standing position without shoes. BMI is calculated using the Quetelet index: weight(kg)/(height(cm))^2^. Waist hip ratio is the ratio of waist circumference (cm) over hip circumference (cm), and waist height ratio is the ratio of waist circumference (cm) over height (cm).

We measured 11 body composition features using whole-body DXA (Discovery A, Hologic Inc., Bedford, MA,USA): whole body lean mass in kilogram (kg), whole body fat in kg and percentage (%) of whole body weight, trunk fat in kg and % of trunk weight, android fat in kg and % of android weight, genoid fat in kg and % of genoid weight, leg fat in kg and percentage of leg weight. In addition, we calculated 6 ratios: trunk fat over whole body fat ratio, android fat over whole body fat ratio, gynoid fat over whole body fat ratio, leg fat over whole body fat ratio, android fat over gynoid fat ratio, whole body fat over whole body lean mass ratio (Supporting information, Figure S1).

A detailed description of the DXA measurement and an indication of the trunk, android and genoid body regions can be found in the Supporting Information.

### 2.3 Metabolic Biomarker Profiling

Blood collection took place between 8 and 9 am after at least 10 hours of fasting. Metabolic biomarkers were quantified from serum samples of 164 individuals using high-throughput ^1^H-NMR metabolomics (Nightingale Health Ltd, Helsinki, Finland). Details of the experimentation and applications of the NMR metabolomics platform have been described previously^[21]^. This method provides simultaneous quantification of routine lipids, lipoprotein subclass profiling with lipid concentrations within 14 subclasses, fatty acid composition, and various low-molecular metabolites including amino acids, ketone bodies and gluconeogenesis-related metabolites in molar concentration units. The 14 lipoprotein subclass sizes were defined as follows: extremely large VLDL with particle diameters from 75 nm upwards and a possible contribution of chylomicrons, five VLDL subclasses (average particle diameters of 64.0 nm, 53.6 nm, 44.5 nm, 36.8 nm, and 31.3 nm), IDL (28.6 nm), three LDL subclasses (25.5 nm, 23.0 nm, and 18.7 nm), and four HDL subclasses (14.3 nm, 12.1 nm, 10.9 nm, and 8.7 nm). The mean size for VLDL, LDL and HDL particles was calculated by weighting the corresponding subclass diameters with their particle concentrations.

Due to the high correlation among the metabolic biomarkers, we only analyzed the 65 biomarkers that have previously been explored for cardiovascular risk by Würtz *et al.*^[22]^ to enhance interpretability. The selection of these biomarkers was based on previous studies using this platform and the current list comprises the total lipid concentrations, fatty acid composition, and low-molecular-weight metabolites, including amino acids, glycolysis-related metabolites, ketone bodies and metabolites involved in fluid balance and immunity (Table S2).

### 2.4 Statistical analysis

For the following metabolic biomarkers, serum levels were below the detection level for at least one measurement: lipid concentration in Extremely Large VLDL (2.3%), Very Large VLDL (3.0%), Large VLDL (1.5%), and Large HDL (2.3%), and these values were considered as missing (Table S2 All metabolic biomarkers were LN-transformed and consecutively Z-scaled (resulting in a mean of 0 and a standard deviation (SD) of 1). To be able to compare the effects of body composition parameters, all measurement levels were Z-scaled.

Partial correlation of metabolic biomarkers and body composition parameters at baseline was determined using a linear mixed model adjusted for age, gender, status (longevity family member or control), lipid lowering medication, hypertension medication (fixed effects) and household (random effect) with the body composition parameters as outcome. A random effect for household was included to account for the potentially increased similarity among household members (85% belong to a couple sharing a household, i.e. 56 couples in our study), as they generally share diet and other lifestyle factors.

To determine the partial correlation of the change in the metabolic biomarker levels and the change in the body composition parameters after the intervention, a linear mixed model was used with the metabolic biomarker levels as outcome and body composition as determinant adjusted for age, gender, status (longevity family member or control), lipid lowering medication, hypertension medication (fixed effects), household, and individual (random effects). For additional analyses, weight was added to the model to determine general weight loss-independent effects.

All statistical analyses were performed with STATA/SE 13.1 and heatmaps were generated using the *heatmap.2* function of the *gplots* package in R. Since we tested 65 metabolic biomarkers and 22 body composition phenotypes we considered p < 3.5 x 10^-5^ (0.05/(65 x 22)) as significant after adjustment for multiple testing.

## 3. Results

### 3.1 Study population

The current investigation of the relation between changing body composition and circulating metabolic biomarkers was performed in a representative subgroup of 132 participants of the GOTO study of whom body composition measures and ^1^H-NMR circulating metabolic biomarkers were available before and after the intervention (Table 1). The mean age of the study participants was 63 years (range 46-75 years), they had a mean BMI of 27 kg/m^2^ (SD 2.4) and 18 participants (11%) were obese (BMI>30 ks/m^2^).

### 3.2 Abdominal fat associates with circulating metabolic biomarkers and body composition on baseline

In order to compare our findings with those from a previous study in younger individuals^[5]^, we first investigated the association between body composition parameters and circulating metabolic biomarkers of the GOTO study at baseline (Figure S2). We confirmed that body composition features, especially large amount of abdominal fat measures, mainly associated with smaller HDL diameter, higher VLDL particle concentrations and higher circulating levels of triglyceride and glycoprotein acetyls (Figure S2). In contrast, in the GOTO study that consisted of older adults, a larger amount of abdominal fat mass was additionally associated with higher circulating concentrations of glycerol and 3-hydroxybutyrate. Furthermore, we observed stronger associations with the DXA fat measures than with the anthropometrics parameters of body composition.

We subsequently investigated whether the associations between body composition and circulating metabolic biomarker levels would still hold after adjustment for BMI. Figure 1 shows a heatmap of the partial correlation (adjusted for BMI) between metabolic biomarkers and body composition parameters at baseline. The hierarchical clustering on basis of the partial correlations between body composition measures and metabolic biomarkers, clusters body composition parameters roughly into 5 clusters (Figure 1, colours at left): 1) Green: DXA measures for abdominal fat, 2) Violet: Anthropometric measures of abdominal fat, 3) Orange: Whole body composition, 4) Yellow: Lower body fat and lean mass, 5) Blue: DXA measures of the ratio between lower body fat and whole body fat. After adjustment for BMI just the DXA measures of abdominal fat and the inversely correlated ratio of lower body fat over whole body fat were associated with circulating metabolic biomarkers. Since the lower body fat measures themselves do not show any association with metabolic biomarkers, the latter association seems to be driven by the whole body fat measure. Hence, after adjustment for BMI, we observed that a higher percentage of fat in the trunk or android body regions (abdominal fat) associated with a lower concentration of lipids in (extra) large HDL particles and smaller HDL diameter, a higher concentration of lipids in (extra) large VLDL particles, and higher circulating levels of leucin, isoleucine, serum triglycerides, glycoprotein acetyls, 3-hydroxybutyrate, and glycerol.

**Figure 1:**
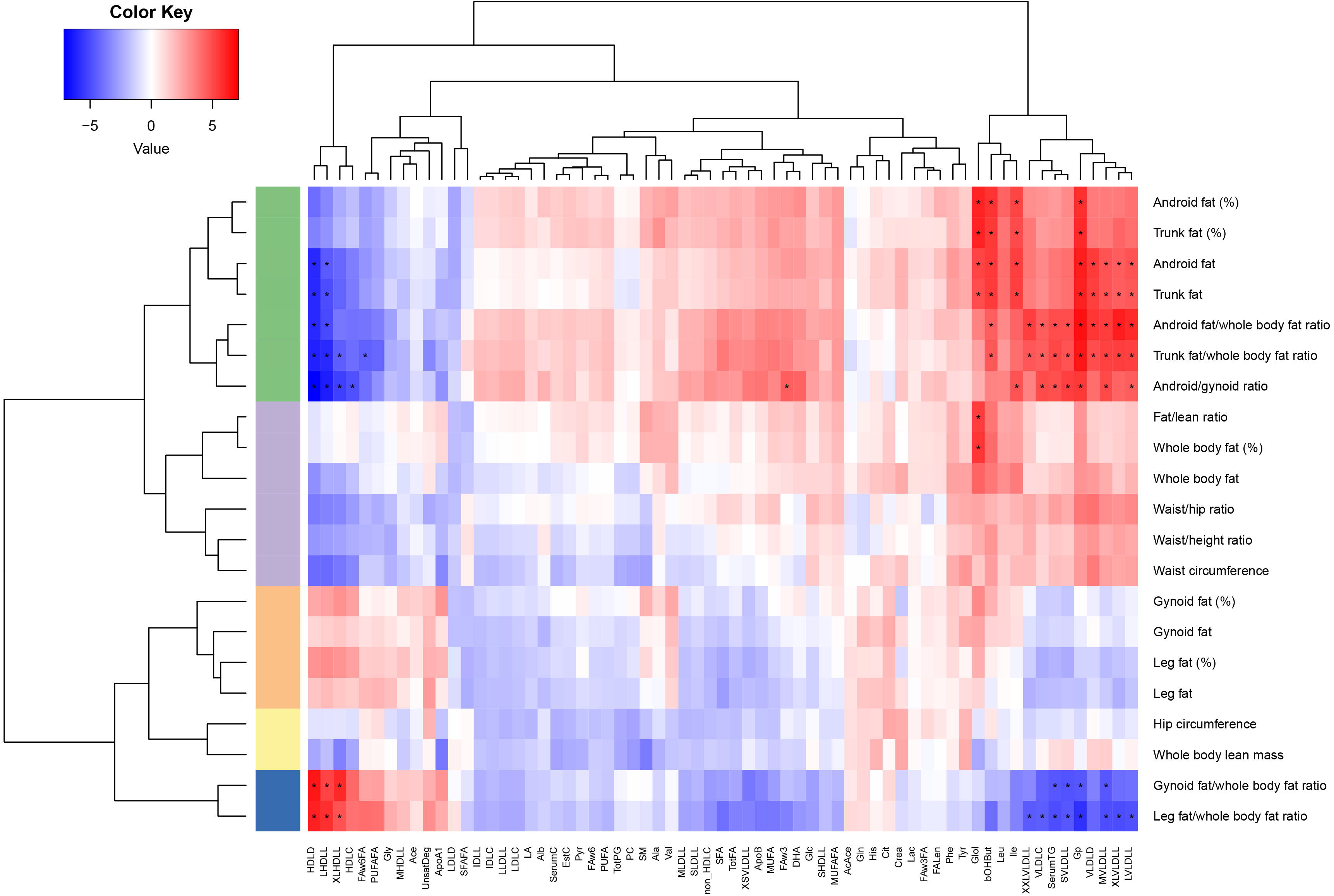
BMI adjusted partial correlation coefficients between circulating metabolic biomarkers and body composition measures at baseline. Android fat (%), gynoid fat (%), trunk fat (%), leg fat (%), whole body fat (%), indicate the ratio of fat mass to total mass in that body area. Fat/lean ratio indicates the ratio of whole body fat mass to whole body lean mass. The blue/red colour key denotes the magnitude of the correlation coefficients. The row colours indicate the clusters of body composition parameters based on their correlations with circulating metabolic biomarkers. Green: abdominal fat; violet: whole body fat; orange: ratio of lower body fat to whole body fat; yellow: lower body fat; blue: lean mass. All metabolic biomarkers were LN transformed and standard normal-transformed. Complete names of the metabolic biomarkers are written down in Table S2. *p < 3.5 x 10^-5^ (0.05/(65 metabolic biomarkers x 22 body composition parameters).

### 3.3 Lifestyle change reduces CVD-associated metabolic biomarkers and abdominal fat

Second, we investigated which of the 65 circulating metabolic biomarkers changed in response to the intervention. In total 46 metabolic biomarkers changed significantly due to the intervention and the most prominent effects were observed for LDL and VLDL subclass concentrations, and levels of apoB, monounsaturated fatty acids, triglycerides and cholesterol (Table S3). The metabolic biomarkers responding to the lifestyle intervention combining less caloric intake and more physical activity, clearly responded in the direction of lower risk for cardiovascular disease as reported by Würtz et al^[6]^.

We also investigated which DXA measure of body composition changed in response to the lifestyle intervention. Both men and women reduced their whole body fat with 1.5% (IQR= −0.5%—2.6%) (Figure 2). As expected, android fat and trunk fat reduced most in both women (−2.4% (IQR= −0.5% Ò −4.7%) and −2.1% (IQR=-0.6% -- 3.4%), respectively) and men (−2.9% (IQR=−0.9% - -5.0%) and −2.3% (IQR= −1.0% −3.6%), respectively). Likewise, waist circumference, waist/hip ratio and fat/lean ratio decreased similarly in men and women (Table S4).

**Figure 2:**
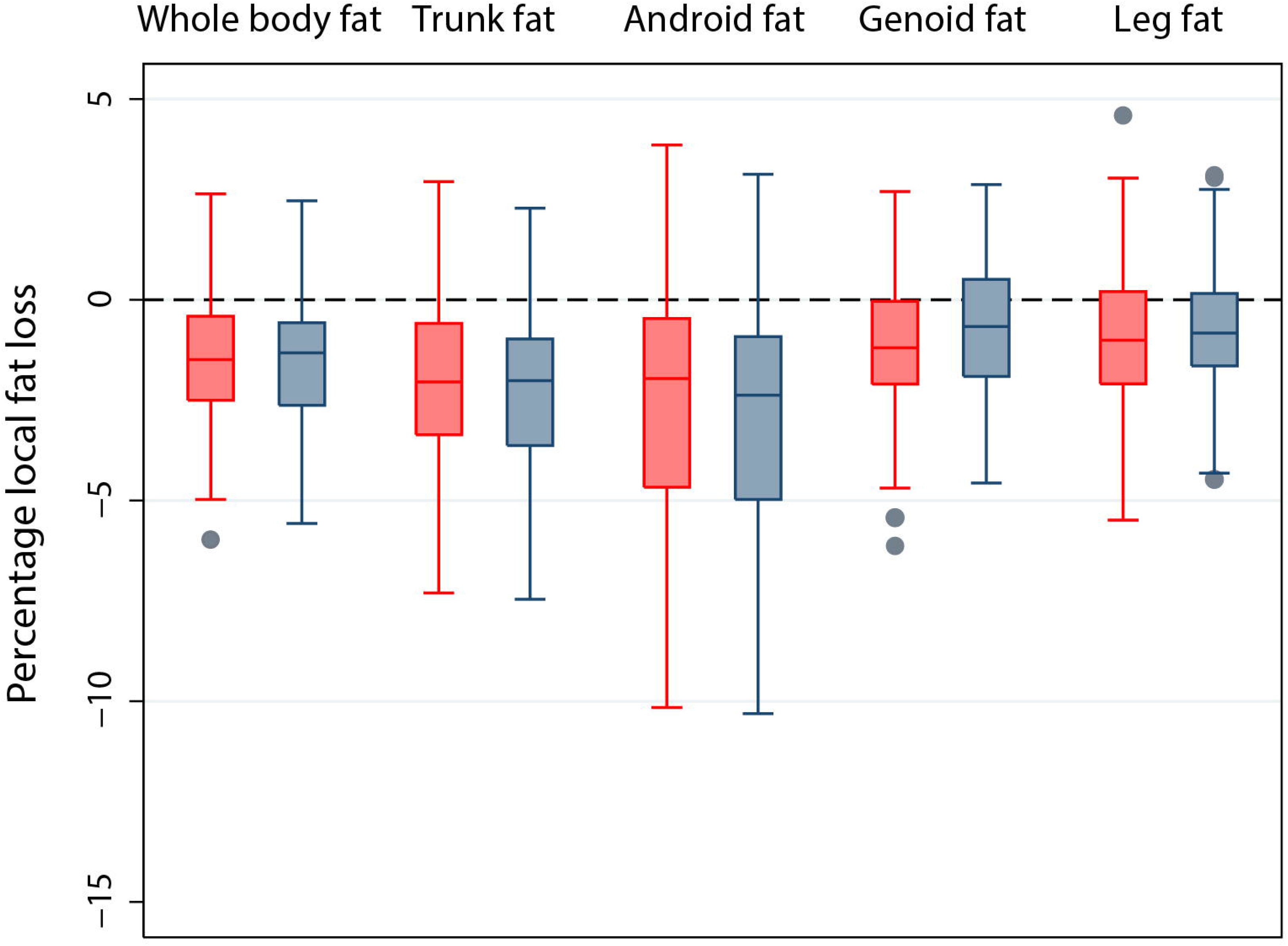
Change in local fat percentage in the whole body, trunk, android, gynoid and leg area after the intervention, stratified for gender. Red: women; Gray: men. The lower and upper boundary of the boxes indicates the interquartile distance (IQR) (the 25^th^ and 75^th^ percentile). The line within the box is the median. The lower whisker indicates the lower adjacent value; the upper whisker indicates the higher adjacent value. Gray dots are individual outliers indicating values that are more than 1.5 times the IQR.

### 3.4 Partial correlations between Δ metabolic biomarkers and Δ abdominal fat

To determine whether the change in circulating metabolic biomarker levels can be explained by the reduction in abdominal fat, we investigated whether the change in abdominal fat (Δ abdominal fat) correlates with the change in the levels of 46 metabolic biomarkers (Δ metabolic biomarker) that altered by the lifestyle intervention (Table S3), while adjusting for general weight loss. The reduction of android and trunk fat, both measured in grams and the percentage of fat in android and trunk, was most strongly associated, independent of general weight loss, with a decrease in circulating glycerol levels (Figure S3). If the android fat mass decreased with 1 SD, glycerol levels decreased with 0. 35 SD (p-value= 1.45 x 10^-11^: Table 2a). Decreasing ratios of abdominal fat over whole body fat, or android fat over genoid fat were most strongly associated with an increasing HDL diameter. If the trunk fat over whole body mass ratio decreased with 1 SD, the HDL diameter increased with 0.36 SD (p-value= 2.56 x 10^-9^: Table 2b).

**Table 2a.** Effect of change in android fat mass on change circulating glycerol levels due to the lifestyle intervention.

| **Glycerol levels\*** | **Effect size** | **CI** | **P-value** |
| --- | --- | --- | --- |
| **Android Fat mass<sup>#</sup>** | **0.35** | **(0.25 - 0.46)** | **1.45 x 10<sup>-11</sup>** |
| Weight (kg) | -0.01 | (-0.03 - 0.00) | 0.024 |
| Age (years) | 0.01 | (0.00 - 0.03) | 0.084 |
| Sex<sup>^</sup> | -0.03 | (-0.23 - 0.16) | 0.736 |
| Status<sup>¥</sup> | 0.04 | (-0.10 - 0.17) | 0.576 |
| Use of lipid lowering medication | 0.01 | (-0.18 - 0.21) | 0.884 |

**Table 2b.** Effect of change in trunk fat over whole body fat ratio on change in HDL diameter due to the lifestyle intervention.

| **HDL diameter\*** | **Effect size** | **CI** | **P-value** |
| --- | --- | --- | --- |
| **Trunk fat/whole body fat ratio<sup>#</sup>** | **-0.36** | **(-0.48 - -0.24)** | **2.56 x 10<sup>-9</sup>** |
| Weight (kg) | -0.03 | (-0.04 - -0.01) | 7.14 x 10<sup>-5</sup> |
| Age (years) | 0.00 | (-0.02 - 0.02) | 0.990 |
| Sex<sup>^</sup> | -0.21 | (-0.45 - 0.03) | 0.093 |
| Status<sup>¥</sup> | 0.27 | (0.06 - 0.49) | 0.013 |
| Use of lipid lowering medication | -0.34 | (-0.65 - -0.03) | 0.029 |

Random effects (mixed) models with glycerol levels(a) or HDL diameter (b) as outcome and android fat mass (a) or trunk fat over whole body mass ratio (b) as determinant, and weight as time-varying covariate, and age, sex, status and lipid medication use as time-invariable covariates.
\* Ln-transformed and Z-scaled
<sup>#</sup> Z-scaled
<sup>^</sup> 0=Female, 1=Male
<sup>¥</sup> 0=Member of long-lived family, 1=Control
CI=Confidence Interval
N=131 individuals contributed to these analyses

In addition, although to a lower extent, the levels of lipids in VLDL particles, serum triglycerides, glycoprotein acetyls, apolipoprotein B, total fatty acids, monounsaturated fatty acids and leucine decreased when abdominal fat reduced.

## 4. Discussion

We investigated the relation between lifestyle intervention-induced changes in body composition, specifically abdominal fat, and the accompanying molecular changes in older adults independent of general weight loss. In young and old individuals abdominal fat mainly associates with a smaller HDL diameter, higher VLDL particle concentrations and higher circulating levels of triglycerides and glycoprotein acetyls. In older adults abdominal fat is additionally associated with higher circulating levels of glycerol and 3-hydroxybutyrate. Furthermore, we showed that after adjustment for BMI, as a measure of overall adiposity, circulating metabolic biomarkers were still associated with abdominal fat. The more abdominal fat, the smaller the HDL diameter, and the lower the concentration of lipids is in HDL particles, and the higher the concentration of lipids in VLDL particles. In addition, if there was more abdominal fat, the circulating levels of glycerol, 3-hydroxybutyrate, leucine and glycoprotein acetyls were higher. This metabolic biomarker profile associated with abdominal fat indicates a high risk for cardiovascular disease^[6]^. We next showed that the intervention beneficially affected especially abdominal fat as well as the majority of the tested metabolic biomarkers (46), of which 26 are known to associate with cardiovascular disease^[6]^. Next, we show that the lifestyle-induced decrease of circulating glycerol levels and increase in HDL diameter can be explained by the loss of abdominal fat. Hence, the lifestyle-induced reduction of abdominal fat in older adults is reflected by decreased circulating glycerol levels and larger HDL diameter.

In older people, measures of BMI or body weight are not able to discriminate with high cardiometabolic disease risk, i.e. low muscle mass and high abdominal fat mass, from those with low risk, i. e. high muscle mass and low abdominal fat mass^[23–25]^. Of the people with an average BMI, around 50% has a percentage of body fat that is too high for their age and gender^[26]^ and 30% is metabolically unhealthy^[27]^. It is known that high body fat and not so much high body weight is associated with an increased risk for cardio-metabolic disease^[28]^. The relatively simple DXA measures for (abdominal) fat mass would then better be able to identify people with high cardiometabolic risk. Because a lifestyle change that reduces caloric intake and increases physical activity may not be beneficial for each older person, it would be crucial to monitor the metabolic effects of lifestyle interventions aimed at reducing the cardio-metabolic disease risk. We found that the healthy reduction in abdominal fat during a lifestyle change was, independent of general weight loss, reflected in by lower circulating glycerol concentrations and larger HDL diameter. Hence, these ^1^H-NMR measures may be further explored to monitor the beneficial effects of a lifestyle change in older people. Circulating glycerol levels and HDL diameter may be valuable tools to monitor cardiometabolic health in older people performing a lifestyle change.

The association of circulating metabolic biomarkers with abdominal fat has been frequently observed in previous studies^[5,29,30]^. We now showed in older adults, that after adjusting for BMI, glycerol particularly associated with abdominal fat measures. Glycerol is produced by white adipose tissue to dispose of excess glucose^[31]^ leading, via hepatic gluconeogenesis, to an increase in circulating glucose levels. A high level of circulating glycerol is a known biomarker for an increasing risk for hyperglycemia and type 2 diabetes^[11]^. Increased HDL diameter also reflects reduced abdominal fat independent of BMI, which is in concordance with previous observations^[32]^. Small, dense HDL subfractions promote cholesterol efflux from foam cell macrophages in the artery wall^[33]^, which would reduce atherosclerotic lesions. We hypothesize that when there is a large amount of abdominal fat, high levels of cholesterol require large cholesterol efflux to clear the foam cells. Hence, when abdominal fat is reduced, for example by a lifestyle intervention, the cholesterol efflux is lowered and the number of small HDL particles is reduced, resulting in higher overall HDL diameter. This suggests, in combination with our findings that older people during a lifestyle intervention mainly loose abdominal fat and decrease their cardiometabolic disease risk, that cardiometabolic disease risk is influenced by abdominal fat, independent of BMI and general weight loss, via circulating glycerol levels and HDL diameter.

The design of the GOTO study has some limitations. The change in lifestyle was for example not controlled, but guided to be feasible for participants. The way participants decreased their caloric intake was different for each participant, which was also the case for the increased physical activity. Though we endeavoured a reduction of 12.5% of caloric intake and 12.5% more physical activity, we are currently not able to analyse to which extent the participants decreased their caloric intake and to which extent they increased their physical activity. Hence, it is unclear whether the changes in metabolic biomarker levels and abdominal fat were mainly due to the change in dietary pattern, physical exercise or the combination of both. This implies that the changes in metabolic biomarkers may be caused by the reduction in abdominal fat due to the lifestyle change, although they may also results from the changes in dietary pattern. Another limitation is that the sample size of the GOTO study does not allow for gender stratified analyses. However, the majority of the female GOTO participants is postmenopausal and sex difference in body composition may therefore be limited. Since body composition and the accompanying cardiometabolic disease risk is a serious issue among older people, the older age of the GOTO study participants is advantageous.

In conclusion, the reduction of abdominal fat in older people due to a lifestyle change, is specifically reflected by decreased circulating glycerol concentration and larger HDL particle diameter, independent of general weight loss. Hence, to monitor the beneficial effects of a lifestyle change at older age circulating glycerol concentration and HDL diameter may be valuable tools.

## Acknowledgements

The research leading to these results has received funding from the European Union’s Seventh Framework Programme (FP7/2007–2011) under grant agreement number 259679. This study was financially supported by the Netherlands Consortium for Healthy Ageing (grant 050-060-810), in the framework of the Netherlands Genomics Initiative, Netherlands Organization for Scientific Research (NWO); by BBMRI-NL, a Research Infrastructure financed by the Dutch government (NWO 184.021.007, 184.033.111) and by the Netherlands CardioVascular Research Initiative (CVON201-03). J. Deelen was financially supported by the Alexander von Humboldt Foundation. The funding agencies had no role in the design and conduct of the study; collection, management, analysis, and interpretation of the data; and preparation, review, or approval of the manuscript.

M.B., B.A.M.S., E.B.A., and P.E.S. designed the study; B.A.M.S., P.D.-S., L.F.G-O., J.G., J.D., O.R., M.A.-K., and D.H. were involved in data acquisition; M.B., B.A.M.S., E.B.A., R.N., and P.E.S. analyzed and interpreted the data; M.B., B.A.M.S., E.B.A., and P.E.S. drafted the manuscript; Critical revision was performed by R.N., J.D., O.R., E.J.M.F., P.E.S.. All the authors read and approved the final manuscript.

We thank all staff members and Bachelor’s and Master’s students who contributed to the preparation, design, and performance of this intervention trial and/or assisted on the project. Last but not least, our gratitude goes to all participants who did their very best to adhere to the intervention guidelines and underwent all measurements.

## Abbreviations

^1^H-NMR: Proton Nuclear Magnetic Resonance
ApoB: Apolipoprotein B
BCAA: Branched-Chain Amino Acid
BMI: Body Mass Index
CVD: Cardiovascular disease
DXA: Dual X-ray Absorptiometry
Glol: Glycerol
GOTO: Growing Old TOgether (Lifestyle intervention study)
HDL: High Density Lipoprotein
LDL: Low Density Lipoprotein
MUFA: Monounsaturated Fatty Acid
SD: Standard deviation
VLDL: Very Low Density Lipoprotein

