## Supplemental Methods and Figures for "Lifestyle-intervention-induced reduction of abdominal fat is reflected by a decreased circulating glycerol level and an increased HDL diameter"

### Supporting methods and figures

| **Page 2** | **Methods DXA measures of body composition  Figure S1.** **Images and recognized body regions from DXA scanner** |
| --- | --- |
| **Page 3** | **Figure S2. Correlation coefficients between serum metabolite levels and body composition measures at baseline without adjustment for BMI** |
| **Page 4** | **Figure S3. Weight loss adjusted correlation coefficients between Δ metabolic biomarkers and Δ abdominal fat measures** |

### Methods Dxa measurement for body composition

Dual energy X-ray absorption (DXA) is used as reliable technology for measuring bone density for the assessment of bone mineral density (BMD) in the hip and the lower spine. In addition, the DXA device is capable of measuring body composition. This technique is based on the difference in attenuation of bone, fat and lean tissue. The X-ray source generates a beam of X-rays with two different energy levels (140 KeV and 100 KeV). During the passage through tissues a part of the X-rays are attenuated. The attenuation is influenced by the intensity of the X-ray energy, and the thickness and density of the tissue. The difference in attenuation of the two X-ray energy peaks is specific for each tissue. A distinction can be made between total body mass, lean mass, fat mass, android fat and genoid fat, using software provided by Hologic Inc (Bedford, MA,USA).

DXA was performed in the supine position with the patient undressed (except underwear) and without any external metal and objects (zipper, bra, piercings) to avoid artifacts and inaccuracy. There were no positioning aids or other objects in the field of view. The patient was positioned on the table with the legs tied together, reducing patient movement during the scan. The arms were positioned along the body with space between the arms and the trunk. If patients did not fit into the field of view, the positioning was modified by standard adjustments. When the patient was taller than 195.6.cm, the feet were positioned outside the field of view and when the width of the patients exceeded 67cm, the left arm was positioned outside the scan field. During post processing, eleven regions of interest (ROI) were drawn to separate the head, upper limb (left and right), lower limb (left and right), thoracic spine, lumbar spine, ribs (left and right), pelvis and trunk (calculated). The android and gynoid fat regions were automatically drawn by the software.

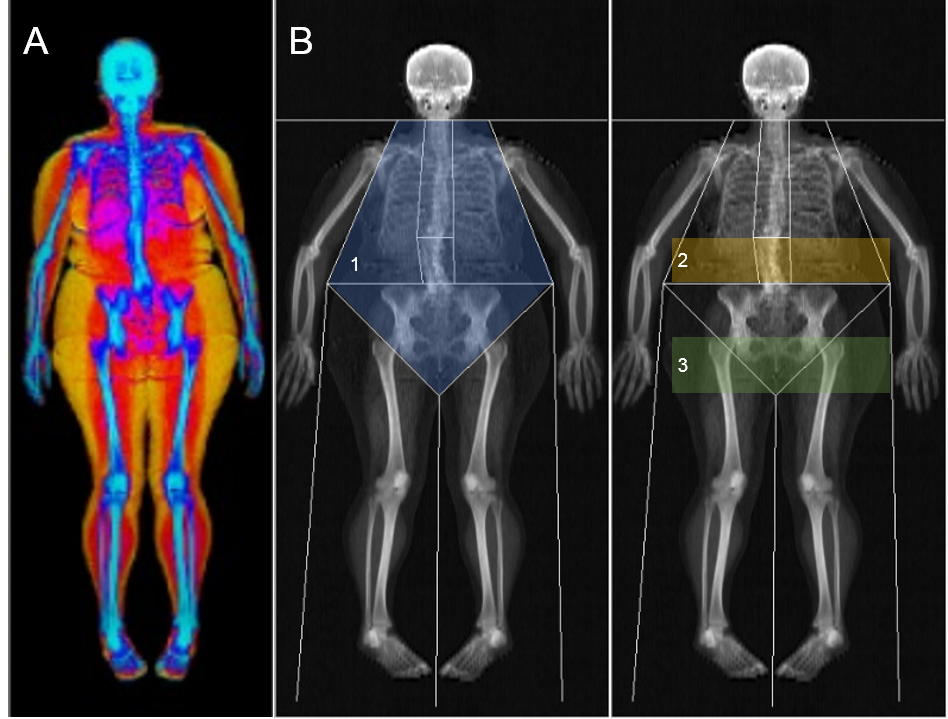

**Figure S1. Images and recognized body regions from DXA scanner. A:** separation between bone (blue), lean (red) and fat (orange). **B:** Indications of body regions of interest are 1) Trunk (blue), 2) Android (orange), Genoid (green).

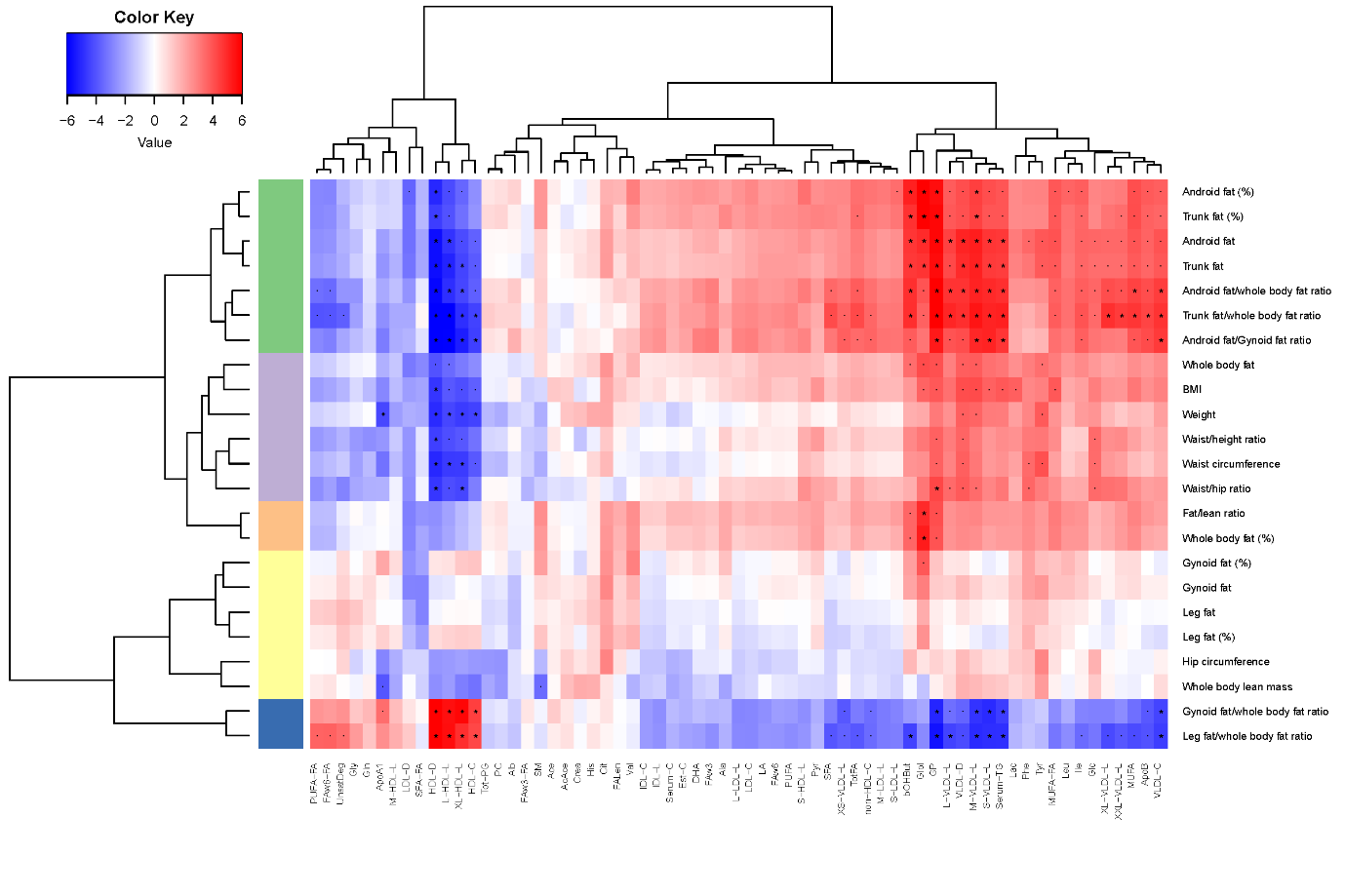

**Figure S2. Correlation coefficients between serum metabolite levels and body composition measures at baseline without adjustment for BMI.** In the hierarchical clustered heatmap the T-statistics for the association between the body composition parameter and metabolite levels was plotted resulting from a linear mixed model with the body composition parameters as outcome and adjusted for age, gender, status (longevity family member or control), lipid lowering medication, hypertension medication (fixed effects) and household (random effects). A random effect for household was included to account for the potentially increased similarity among household members (56 couples in our study), as they generally share diet and other lifestyle factors. Android fat (%), gynoid fat (%), trunk fat (%), leg fat (%), whole body fat (%), indicate the ratio of fat mass to total mass in that body area. Fat/lean ratio indicates the ratio of whole body fat mass to whole body lean mass. The colour key denotes the magnitude of the correlation coefficients. The row colours indicate the clusters of body composition parameters based on their correlations with serum metabolite levels. Green: abdominal fat; violet: anthropometrics; orange: whole body composition; yellow: lower body fat and lean mass; blue: ratio of lower body fat to whole body fat. All metabolite levels were LN transformed and standard normal-transformed. Complete names of the metabolites are written down in Table S2. The P-values denote the statistical significance after correcting for multiple testing: P-values *p < 3.5 x 10^-5^; - p < 0.001.

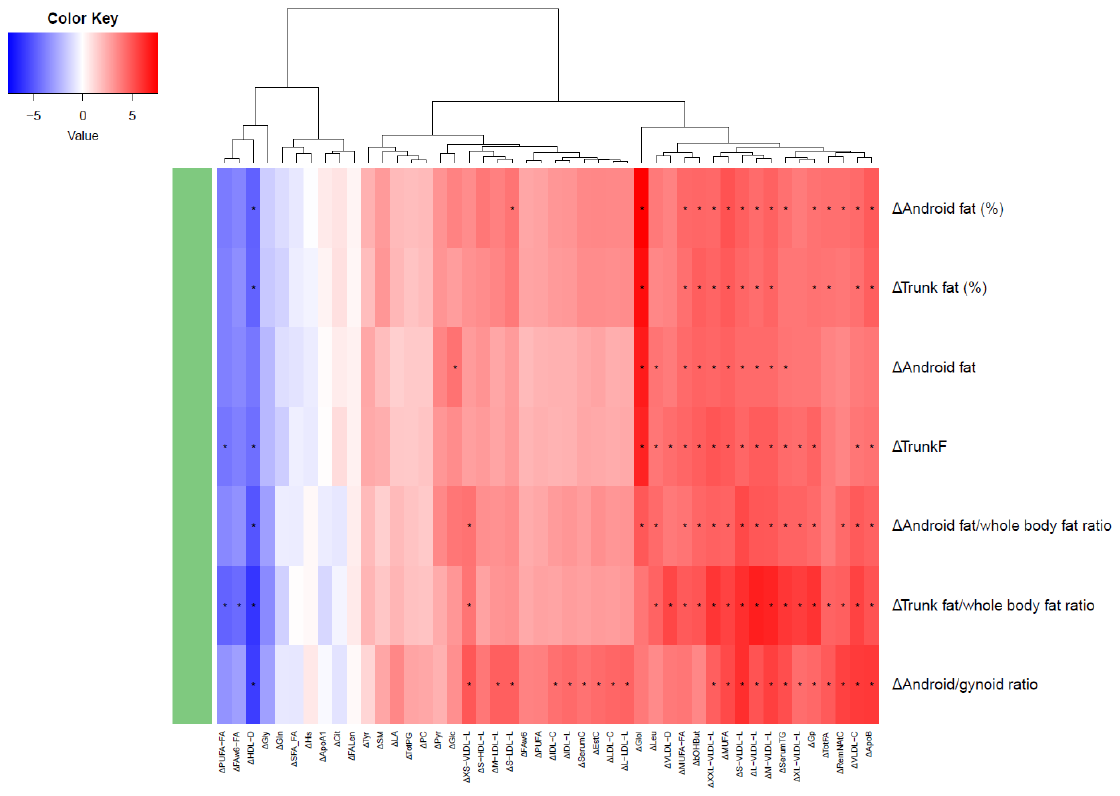

**Figure S3: Weight loss adjusted partial correlation coefficients between Δ metabolic biomarkers and Δ abdominal fat measures.**

Android fat (%), Gynoid fat (%), Trunk fat (%), Leg fat (%), Whole body fat (%), indicate the ratio of fat mass to total tissue mass in that body area. The color key denotes the magnitude of the t-statistics of the regression between the change (Δ) in metabolite levels and the change (Δ) in body composition parameter. All metabolite levels were LN transformed and standard normal-transformed. Complete names of the metabolites are written down in Table S2. The P-values denote the statistical significance after Bonferroni correcting for multiple testing: P-values *p < 0.00015. In the hierarchical clustered heatmap the T-statistics for the association between the Δmetabolite levels and Δbody composition parameter was plotted resulting from a linear mixed model with the Δmetabolite levels as outcome and Δbody composition parameter as determinant adjusted for weight, age, gender, status (longevity family member or control), lipid lowering medication, hypertension medication (fixed effects) and household (random effects). A random effect for household was included to account for the potentially increased similarity among household members (56 couples in our study), as they generally share diet and other lifestyle factors, and another random effect for individuals to take into account the timepoints at baseline and after the intervention.
