## Supplemental Tables for "Lifestyle-intervention-induced reduction of abdominal fat is reflected by a decreased circulating glycerol level and an increased HDL diameter"

**Table S1. Baseline characteristics of the complete GOTO study population***

| | N | Mean | SD |
|---|---|---|---|
| Age (years) | 164 | 63.0 | 5.7 |
| % Female | 81 | 49.4 | |
| % Lipid lowering medication | 29 | 17.7 | |
| % Antihypertensive medication | 51 | 31.1 | |
| **Anthropometrics** | | | |
| Height (m) | 163 | 1.7 | 0.1 |
| Weight (kg) | 163 | 79.4 | 9.8 |
| BMI (kg/m2) | 163 | 27.0 | 2.5 |
| Waist circumference (cm) | 164 | 96.1 | 8.0 |
| Hip circumference (cm) | 164 | 104.3 | 5.4 |
| Waist/hip ratio (cm) | 164 | 0.9 | 0.1 |
| Waist/height ratio | 163 | 56.1 | 4.6 |
| **DXA measures** | | | |
| Whole body lean mass (kg) | 142 | 54.1 | 9.7 |
| Whole body fat (kg) | 142 | 25.6 | 6.2 |
| Whole body fat (%) | 142 | 32.3 | 7.3 |
| Trunk fat (kg) | 142 | 13.1 | 3.6 |
| Trunk fat (%) | 142 | 32.7 | 7.2 |
| Android fat (kg) | 142 | 2.2 | 0.7 |
| Android fat (%) | 142 | 35.1 | 7.3 |
| Gynoid fat (kg) | 142 | 4.1 | 1.1 |
| Gynoid fat (%) | 142 | 32.8 | 8.2 |
| Leg fat (kg) | 142 | 8.32 | 2.67 |
| Leg fat (%) | 142 | 32.13 | 9.36 |
| Trunk fat /whole body fat ratio | 142 | 0.51 | 0.06 |
| Adroid fat /whole body fat ratio | 142 | 0.09 | 0.02 |
| Gynoid fat /whole body fat ratio | 142 | 0.16 | 0.02 |
| Leg fat /whole body fat ratio | 142 | 0.32 | 0.06 |
| Android fat/gynoid fat ratio | 142 | 1.10 | 0.20 |
| Whole body fat/Whole body lean mass ratio* | 142 | 0.49 | 0.17 |

*The complete group 164 participants
SD= standard deviation

**Table S2. Nomenclature of the 65 metabolic biomarkers**

| Parameter | Description | Percentage missing values |
|---|---|---|
| XXL-VLDL-L | Total lipids in chylomicrons and extremely large VLDL (mmol/l) | 2.3 |
| XL-VLDL-L | Total lipids in very large VLDL (mmol/l) | 3.0 |
| L-VLDL-L | Total lipids in large VLDL (mmol/l) | 1.5 |
| M-VLDL-L | Total lipids in medium VLDL (mmol/l) | 0.0 |
| S-VLDL-L | Total lipids in small VLDL (mmol/l) | 0.0 |
| XS-VLDL-L | Total lipids in very small VLDL (mmol/l) | 0.0 |
| IDL-L | Total lipids in IDL (mmol/l) | 0.0 |
| L-LDL-L | Total lipids in large LDL (mmol/l) | 0.0 |
| M-LDL-L | Total lipids in medium LDL (mmol/l) | 0.0 |
| S-LDL-L | Total lipids in small LDL (mmol/l) | 0.0 |
| XL-HDL-L | Total lipids in very large HDL (mmol/l) | 0.0 |
| L-HDL-L | Total lipids in large HDL (mmol/l) | 2.3 |
| M-HDL-L | Total lipids in medium HDL (mmol/l) | 0.0 |
| S-HDL-L | Total lipids in small HDL (mmol/l) | 0.0 |
| VLDL-D | Mean diameter for VLDL particles (nm) | 0.0 |
| LDL-D | Mean diameter for LDL particles (nm) | 0.0 |
| HDL-D | Mean diameter for HDL particles (nm) | 0.0 |
| Serum_C | Serum total cholesterol (mmol/l) | 0.0 |
| non-HDL-C | Total cholesterol in non-HDL (mmol/l) | 0.0 |
| VLDL-C | Total cholesterol in VLDL (mmol/l) | 0.0 |
| IDL-C | Total cholesterol in IDL (mmol/l) | 0.0 |
| LDL-C | Total cholesterol in LDL (mmol/l) | 0.0 |
| HDL-C | Total cholesterol in HDL (mmol/l) | 0.0 |
| Est-C | Esterified cholesterol (mmol/l) | 1.5 |
| Serum-TG | Serum total triglycerides (mmol/l) | 0.0 |
| Tot-PG | Total phosphoglycerides (mmol/l) | 1.5 |
| PC | Phosphatidylcholine and other cholines (mmol/l) | 1.5 |
| SM | Sphingomyelins (mmol/l) | 1.5 |
| ApoA1 | Apolipoprotein A-I (g/l) | 0.0 |
| ApoB | Apolipoprotein B (g/l) | 0.0 |
| Tot-FA | Total fatty acids (mmol/l) | 1.5 |
| FALen | Estimated description of fatty acid chain length, not actual carbon number () | 1.5 |
| UnsatDeg | Estimated degree of unsaturation () | 1.5 |
| DHA | 22:6, docosahexaenoic acid (mmol/l) | 1.5 |
| LA | 18:2, linoleic acid (mmol/l) | 1.5 |
| FAw3 | Omega-3 fatty acids (mmol/l) | 1.5 |
| FAw6 | Omega-6 fatty acids (mmol/l) | 1.5 |
| PUFA | Polyunsaturated fatty acids (mmol/l) | 1.5 |
| MUFA | Monounsaturated fatty acids; 16:1, 18:1 (mmol/l) | 1.5 |
| SFA | Saturated fatty acids (mmol/l) | 1.5 |
| FAw3_FA | Ratio of omega-3 fatty acids to total fatty acids (%) | 1.5 |
| FAw6_FA | Ratio of omega-6 fatty acids to total fatty acids (%) | 1.5 |
| PUFA_FA | Ratio of polyunsaturated fatty acids to total fatty acids (%) | 1.5 |
| MUFA_FA | Ratio of monounsaturated fatty acids to total fatty acids (%) | 1.5 |
| SFA_FA | Ratio of saturated fatty acids to total fatty acids (%) | 1.5 |
| Glc | Glucose (mmol/l) | 0.0 |
| Lac | Lactate (mmol/l) | 0.0 |
| Pyr | Pyruvate (mmol/l) | 0.8 |
| Cit | Citrate (mmol/l) | 0.0 |
| Glol | Glycerol (mmol/l) | 1.5 |
| Ala | Alanine (mmol/l) | 0.8 |
| Gln | Glutamine (mmol/l) | 0.0 |
| Gly | Glycine (mmol/l) | 0.0 |
| His | Histidine (mmol/l) | 0.0 |
| Ile | Isoleucine (mmol/l) | 0.8 |
| Leu | Leucine (mmol/l) | 0.8 |
| Val | Valine (mmol/l) | 1.5 |
| Phe | Phenylalanine (mmol/l) | 0.0 |
| Tyr | Tyrosine (mmol/l) | 0.0 |
| Ace | Acetate (mmol/l) | 0.0 |
| AcAce | Acetoacetate (mmol/l) | 0.0 |
| bOHBut | 3-hydroxybutyrate (mmol/l) | 1.5 |
| Crea | Creatinine (mmol/l) | 0.0 |
| Alb | Albumin (signal area) | 0.0 |
| GP | Glycoprotein acetyls, mainly a1-acid glycoprotein (mmol/l) | 0.0 |

**Table S3. Significance of effect of lifestyle change on 1H-NMR metabolic biomarkers**

*According to Supplemental table 2B from Van der Rest et al, Aging 2016 doi: 10.18632/aging.100877*

| Parameter | Description | N | Coefficient* | CI lower bound | CI upper bound | p-value |
|---|---|---|---|---|---|---|
| XXL-VLDL-L | Total lipids in chylomicrons and extremely large VLDL (mmol/l) | 115 | -0.24 | -0.40 | -0.09 | 1.96E-03 |
| XL-VLDL-L | Total lipids in very large VLDL (mmol/l) | 103 | -0.31 | -0.47 | -0.14 | 2.36E-04 |
| L-VLDL-L | Total lipids in large VLDL (mmol/l) | 119 | -0.29 | -0.42 | -0.16 | 1.10E-05 |
| M-VLDL-L | Total lipids in medium VLDL (mmol/l) | 134 | -0.23 | -0.34 | -0.12 | 2.86E-05 |
| S-VLDL-L | Total lipids in small VLDL (mmol/l) | 134 | -0.28 | -0.39 | -0.18 | 2.23E-07 |
| XS-VLDL-L | Total lipids in very small VLDL (mmol/l) | 134 | -0.28 | -0.40 | -0.17 | 1.28E-06 |
| IDL-L | Total lipids in IDL (mmol/l) | 134 | -0.24 | -0.34 | -0.15 | 1.39E-06 |
| L-LDL-L | Total lipids in large LDL (mmol/l) | 134 | -0.23 | -0.33 | -0.13 | 6.53E-06 |
| M-LDL-L | Total lipids in medium LDL (mmol/l) | 134 | -0.24 | -0.35 | -0.13 | 1.32E-05 |
| S-LDL-L | Total lipids in small LDL (mmol/l) | 134 | -0.27 | -0.37 | -0.16 | 7.95E-07 |
| XL-HDL-L | Total lipids in very large HDL (mmol/l) | 133 | 0.07 | -0.05 | 0.19 | 2.53E-01 |
| L-HDL-L | Total lipids in large HDL (mmol/l) | 127 | 0.08 | -0.01 | 0.17 | 6.89E-02 |
| M-HDL-L | Total lipids in medium HDL (mmol/l) | 134 | -0.12 | -0.25 | 0.02 | 8.94E-02 |
| S-HDL-L | Total lipids in small HDL (mmol/l) | 134 | -0.20 | -0.37 | -0.03 | 2.22E-02 |
| VLDL-D | Mean diameter for VLDL particles (nm) | 134 | -0.14 | -0.27 | 0.00 | 4.20E-02 |
| LDL-D | Mean diameter for LDL particles (nm) | 134 | 0.12 | -0.07 | 0.30 | 2.26E-01 |
| HDL-D | Mean diameter for HDL particles (nm) | 134 | 0.11 | 0.01 | 0.21 | 2.86E-02 |
| Serum_C | Serum total cholesterol (mmol/l) | 130 | -0.19 | -0.31 | -0.06 | 2.91E-03 |
| non-HDL-C | Total cholesterol in non-HDL (mmol/l) | 151 | -0.25 | -0.36 | -0.14 | 4.07E-06 |
| VLDL-C | Total cholesterol in VLDL (mmol/l) | 134 | -0.30 | -0.42 | -0.18 | 3.56E-07 |
| IDL-C | Total cholesterol in IDL (mmol/l) | 134 | -0.27 | -0.37 | -0.17 | 1.94E-07 |
| LDL-C | Total cholesterol in LDL (mmol/l) | 134 | -0.25 | -0.35 | -0.14 | 4.68E-06 |
| HDL-C | Total cholesterol in HDL (mmol/l) | 134 | 0.01 | -0.08 | 0.09 | 8.99E-01 |
| Est-C | Esterified cholesterol (mmol/l) | 130 | -0.31 | -0.42 | -0.20 | 1.28E-08 |
| Serum-TG | Serum total triglycerides (mmol/l) | 134 | -0.18 | -0.29 | -0.07 | 1.80E-03 |
| Tot-PG | Total phosphoglycerides (mmol/l) | 130 | -0.26 | -0.39 | -0.13 | 1.30E-04 |
| PC | Phosphatidylcholine and other cholines (mmol/l) | 130 | -0.17 | -0.30 | -0.04 | 8.00E-03 |
| SM | Sphingomyelins (mmol/l) | 130 | -0.26 | -0.36 | -0.16 | 3.73E-07 |
| ApoA1 | Apolipoprotein A-I (g/l) | 134 | -0.15 | -0.25 | -0.05 | 4.21E-03 |
| ApoB | Apolipoprotein B (g/l) | 134 | -0.31 | -0.41 | -0.22 | 1.12E-10 |
| Tot-FA | Total fatty acids (mmol/l) | 130 | -0.22 | -0.34 | -0.10 | 3.87E-04 |
| FALen | Estimated description of fatty acid chain length, not actual carbon number () | 130 | 0.41 | 0.22 | 0.61 | 2.43E-05 |
| UnsatDeg | Estimated degree of unsaturation () | 130 | -0.12 | -0.28 | 0.03 | 1.10E-01 |
| DHA | 22:6, docosahexaenoic acid (mmol/l) | 130 | -0.06 | -0.17 | 0.06 | 3.44E-01 |
| LA | 18:2, linoleic acid (mmol/l) | 130 | -0.26 | -0.38 | -0.14 | 1.75E-05 |
| FAw3 | Omega-3 fatty acids (mmol/l) | 130 | -0.02 | -0.18 | 0.15 | 8.40E-01 |
| FAw6 | Omega-6 fatty acids (mmol/l) | 130 | -0.17 | -0.31 | -0.03 | 1.85E-02 |
| PUFA | Polyunsaturated fatty acids (mmol/l) | 130 | -0.32 | -0.43 | -0.20 | 5.03E-08 |
| MUFA | Monounsaturated fatty acids; 16:1, 18:1 (mmol/l) | 130 | -0.21 | -0.32 | -0.10 | 2.11E-04 |
| SFA | Saturated fatty acids (mmol/l) | 130 | -0.10 | -0.24 | 0.04 | 1.50E-01 |
| FAw3_FA | Ratio of omega-3 fatty acids to total fatty acids (%) | 130 | -0.02 | -0.18 | 0.15 | 8.40E-01 |
| FAw6_FA | Ratio of omega-6 fatty acids to total fatty acids (%) | 130 | -0.17 | -0.31 | -0.03 | 1.85E-02 |
| PUFA_FA | Ratio of polyunsaturated fatty acids to total fatty acids (%) | 130 | -0.17 | -0.31 | -0.02 | 2.24E-02 |
| MUFA_FA | Ratio of monounsaturated fatty acids to total fatty acids (%) | 130 | -0.13 | -0.25 | -0.01 | 2.85E-02 |
| SFA_FA | Ratio of saturated fatty acids to total fatty acids (%) | 130 | 0.38 | 0.19 | 0.57 | 7.53E-05 |
| Glc | Glucose (mmol/l) | 162 | -0.24 | -0.38 | -0.11 | 4.98E-04 |
| Lac | Lactate (mmol/l) | 162 | -0.05 | -0.23 | 0.13 | 6.02E-01 |
| Pyr | Pyruvate (mmol/l) | 161 | -0.26 | -0.43 | -0.09 | 2.26E-03 |
| Cit | Citrate (mmol/l) | 162 | 0.20 | 0.05 | 0.35 | 1.00E-02 |
| Glol | Glycerol (mmol/l) | 157 | -0.17 | -0.30 | -0.04 | 1.08E-02 |
| Ala | Alanine (mmol/l) | 161 | 0.01 | -0.13 | 0.15 | 8.72E-01 |
| Gln | Glutamine (mmol/l) | 162 | 0.19 | 0.06 | 0.33 | 4.43E-03 |
| Gly | Glycine (mmol/l) | 161 | 0.20 | 0.12 | 0.29 | 1.50E-06 |
| His | Histidine (mmol/l) | 162 | -0.52 | -0.71 | -0.34 | 3.23E-08 |
| Ile | Isoleucine (mmol/l) | 161 | -0.12 | -0.24 | 0.00 | 5.30E-02 |
| Leu | Leucine (mmol/l) | 161 | -0.18 | -0.31 | -0.05 | 8.22E-03 |
| Val | Valine (mmol/l) | 160 | -0.12 | -0.26 | 0.03 | 1.08E-01 |
| Phe | Phenylalanine (mmol/l) | 162 | -0.01 | -0.17 | 0.14 | 8.52E-01 |
| Tyr | Tyrosine (mmol/l) | 162 | -0.24 | -0.41 | -0.07 | 5.35E-03 |
| Ace | Acetate (mmol/l) | 162 | -0.01 | -0.13 | 0.11 | 8.48E-01 |
| AcAce | Acetoacetate (mmol/l) | 162 | -0.10 | -0.26 | 0.07 | 2.41E-01 |
| bOHBut | 3-hydroxybutyrate (mmol/l) | 158 | -0.15 | -0.30 | -0.01 | 3.77E-02 |
| Crea | Creatinine (mmol/l) | 162 | -0.09 | -0.19 | 0.01 | 7.10E-02 |
| Alb | Albumin (signal area) | 162 | -0.07 | -0.22 | 0.08 | 3.64E-01 |
| GP | Glycoprotein acetyls, mainly a1-acid glycoprotein (mmol/l) | 162 | -0.15 | -0.27 | -0.04 | 7.83E-03 |

*The significance of the effects of the intervention on metabolite measures were determined using a linear mixed model adjusted for age, gender, status (longevity family member or control) (fixed effects), household, and individual (random effects).

Green p-values indicate nominal significant (p-value<0.05) changes in metabolite levels due to the lifestyle intervention (46 metabolites).

CI= confidence interval

**Table S4. Changes in body composition due to the lifestyle intervention**

| | ALL | | | | FEMALE | | | | MALE | | | |
|---|---|---|---|---|---|---|---|---|---|---|---|---|
| | N | Mean change | SD | p-value* | N | Mean change | SD | p-value# | N | Mean change | SD | p-value# |
| **Anthropometrics** | | | | | | | | | | | | |
| Weight (kg) | 132 | -3.36 | 2.33 | **<0.001** | 65 | -3.35 | 2.09 | **<0.001** | 67 | -3.37 | 2.55 | **<0.001** |
| BMI (kg/m2) | 132 | -1.14 | 0.80 | **<0.001** | 65 | -1.24 | 0.79 | **<0.001** | 67 | -1.05 | 0.81 | **<0.001** |
| Waist circumference (cm) | 132 | -4.51 | 5.44 | **<0.001** | 65 | -4.60 | 5.43 | **<0.001** | 67 | -4.42 | 5.50 | **<0.001** |
| Hip circumference (cm) | 132 | -2.95 | 4.30 | **<0.001** | 65 | -3.53 | 4.29 | **<0.001** | 67 | -2.40 | 4.26 | **<0.001** |
| Waist/hip ratio (cm) | 132 | -0.02 | 0.05 | **<0.001** | 64 | -0.02 | 0.05 | 0.018 | 67 | -0.02 | 0.04 | **<0.001** |
| Waist/height ratio | 132 | -2.65 | 3.23 | **<0.001** | 64 | -2.81 | 3.32 | **<0.001** | 67 | -2.49 | 3.16 | **<0.001** |
| **DXA measures** | | | | | | | | | | | | |
| Whole body lean mass (kg) | 132 | -1.2 | 1.2 | **<0.001** | 65 | -1.1 | 1.2 | **<0.001** | 67 | -1.4 | 1.2 | **<0.001** |
| Whole body fat (kg) | 132 | -2.2 | 1.8 | **<0.001** | 65 | -2.3 | 1.7 | **<0.001** | 67 | -2.1 | 1.9 | **<0.001** |
| Whole body fat (%) | 132 | -1.5 | 1.7 | **<0.001** | 65 | -1.5 | 1.7 | **<0.001** | 67 | -1.5 | 1.7 | **<0.001** |
| Trunk fat (kg) | 132 | -1.5 | 1.1 | **<0.001** | 65 | -1.5 | 1.1 | **<0.001** | 67 | -1.6 | 1.2 | **<0.001** |
| Trunk fat (%) | 132 | -2.2 | 2.2 | **<0.001** | 65 | -2.1 | 2.2 | **<0.001** | 67 | -2.3 | 2.2 | **<0.001** |
| Android fat (kg) | 132 | -0.3 | 0.3 | **<0.001** | 65 | -0.3 | 0.3 | **<0.001** | 67 | -0.4 | 0.3 | **<0.001** |
| Android fat (%) | 132 | -2.6 | 3.0 | **<0.001** | 65 | -2.4 | 3.1 | **<0.001** | 67 | -2.9 | 2.9 | **<0.001** |
| Gynoid fat (kg) | 132 | -0.3 | 0.3 | **<0.001** | 65 | -0.4 | 0.3 | **<0.001** | 67 | -0.2 | 0.3 | **<0.001** |
| Gynoid fat (%) | 132 | -0.9 | 1.8 | **<0.001** | 65 | -1.1 | 1.9 | **<0.001** | 67 | -0.7 | 1.6 | **<0.001** |
| Leg fat (kg) | 132 | -0.5 | 0.7 | **<0.001** | 65 | -0.7 | 0.8 | **<0.001** | 67 | -0.4 | 0.6 | **<0.001** |
| Leg fat (%) | 132 | -0.8 | 1.7 | **<0.001** | 65 | -0.9 | 1.8 | **<0.001** | 67 | -0.7 | 1.6 | **<0.001** |
| Trunk fat /whole body fat ratio | 132 | 0.81 | 1.6 | **<0.001** | 65 | 0.83 | 2.0 | **<0.001** | 67 | 0.78 | 1.0 | **<0.001** |
| Android fat /whole body fat ratio | 132 | 0.23 | 1.1 | **<0.001** | 65 | 0.33 | 1.5 | **<0.001** | 67 | 0.14 | 0.2 | **<0.001** |
| Gynoid fat /whole body fat ratio | 132 | 0.06 | 0.9 | **<0.001** | 65 | 0.12 | 0.8 | **<0.001** | 67 | 0.00 | 1.0 | **<0.001** |
| Leg fat /whole body fat ratio | 132 | 0.11 | 1.6 | **<0.001** | 65 | 0.10 | 2.0 | **<0.001** | 67 | 0.11 | 1.2 | **<0.001** |
| Android fat/gynoid fat ratio | 132 | -0.06 | 0.1 | **<0.001** | 65 | -0.04 | 0.1 | **<0.001** | 67 | -0.09 | 0.1 | **<0.001** |
| Whole body fat/Whole body lean mass ratio | 132 | -0.03 | 0.0 | **<0.001** | 65 | -0.04 | 0.0 | **<0.001** | 67 | -0.03 | 0.0 | **<0.001** |

*The significance of the effects of the intervention on body composition measures were determined using a linear mixed model adjusted for age, gender, status (longevity family member or control) (fixed effects), household, and individual (random effects).

#The significance of the effects of the intervention on body composition measures were determined using a linear mixed model adjusted for age, status (longevity family member or control) (fixed effects), and individual (random effects).

SD= standard deviation
